# Optogenetic control of protein binding using light-switchable nanobodies

**DOI:** 10.1101/739201

**Authors:** Agnieszka A. Gil, Evan M. Zhao, Maxwell Z. Wilson, Alexander G. Goglia, Cesar Carrasco-Lopez, José L. Avalos, Jared E. Toettcher

## Abstract

A growing number of optogenetic tools have been developed to control binding between two engineered protein domains. In contrast, relatively few tools confer light-switchable binding to a generic target protein of interest. Such a capability would offer substantial advantages, enabling photoswitchable binding to endogenous target proteins *in vivo* or light-based protein purification *in vitro*. Here, we report the development of opto-nanobodies (OptoNBs), a versatile class of chimeric photoswitchable proteins whose binding to proteins of interest can be enhanced or inhibited upon blue light illumination. We find that OptoNBs are suitable for a range of applications: modulating intracellular protein localization and signaling pathway activity and controlling target protein binding to surfaces and in protein separation columns. This work represents a first step towards programmable photoswitchable regulation of untagged, endogenous target proteins.

**Highlights:**

1. Opto-Nanobodies (OptoNBs) enable light-regulated binding to a wide range of protein targets.
2. We identify an optimized LOV domain and two loop insertion sites for light-regulated binding.
3. OptoNBs function *in vivo*: when expressed in cells and fused to signaling domains, OptoNBs enable light-activated and light-inhibited Ras/Erk signaling.
4. OptoNBs function *in vitro*: Target proteins can be reversibly bound to OptoNB-coated beads and separated through size-exclusion chromatography.

## Introduction

Nearly 20 years after the initial development of light-regulated transcription in yeast^1^ and light-gated ion channels in neuroscience^2^, optogenetics has been extended to almost every corner of cell biology. Optogenetic proteins are now available to control the fundamental currencies of protein heterodimerization^3–5^, homo-dimerization^6,7^, gene expression^1,8,9^, degradation^10^, nuclear-cytosolic translocation^11–14^, and even liquid-liquid protein phase separation^15,16^. These techniques have enabled a new generation of precise perturbation studies to interrogate how the timing, spatial location, and identity of active proteins alter cellular and developmental processes.

Yet despite this growing suite of optogenetic tools, some applications have remained elusive. Light-gated protein-protein interactions are typically induced between a light-sensitive protein and a target peptide that is derived from the original plant or cyanobacterial host (e.g., dimerization between PhyB/PIF6 or Cry2/CIB)^3,5,6^, or an engineered, synthetic protein-protein interaction (e.g., binding of an engineered AsLOV2 variant to a PDZ or SSPB peptide or between Dronpa monomers)^4,7,17^. In contrast, achieving light-switchable binding to an arbitrary target protein of interest has remained elusive. The ability to reversibly bind and release a protein of interest in response to light would hold considerable promise for reversibly regulating endogenous signaling activity *in vivo*, developing biologics that can be precisely targeted in space and time, and enabling protein purification without affinity tags.

Here, we present opto-nanobodies (OptoNBs): a class of engineered proteins capable of reversible, light-controlled binding against multiple protein targets. Nanobodies, small binding proteins formed from the single variable domain of camelid antibodies, provide a versatile scaffold for obtaining binding to a broad range of target epitopes and are functional in both intracellular and extracellular environments^18^. Our approach for obtaining photoswitchable nanobodies builds on recent pioneering work to insert a photoswitchable light-oxygen-voltage (LOV) domain into solvent-exposed loops on proteins of interest^19^. We identify loop insertion sites and LOV domain variants that trigger a light-inducible change in binding in three nanobodies against two model target proteins, EGFP and mCherry. We further demonstrate that nanobody binding can be applied *in vivo* and *in vitro*, in applications ranging from control over mammalian Ras/Erk signaling to photoswitchable binding on agarose beads and in size exclusion columns. The OptoNB platform opens the door to developing light-switchable binders against a broad range of protein targets and may thus represent a first step toward a new class of photoswitchable biologics.

## Results

### Initial optoNBs exhibit weak light-switchable binding as well as nuclear export

Our strategy to engineer light-controlled nanobodies is based on pioneering work using ligand- or light-gated ‘hairpins’: small domains that can be inserted in-frame into a solvent-exposed loop, whose conformation changes upon illumination or addition of a small molecule^19,20^. We reasoned that by inserting a light-oxygen-voltage sensing domain from *Avena sativa* Phototropin 1 (AsLOV2) into the solvent-exposed loop of a nanobody, it may be possible to allosterically alter the conformation of its binding surface, disrupting recognition of a target protein (Figure 1A). As a starting point, we focused on regulating binding between a model target protein, mCherry, and the LaM8 anti-mCherry nanobody^21^. We took a broad approach to identifying potential AsLOV2 insertion sites, testing all eight conserved, solvent-exposed loops in the nanobody structure excluding the hypervariable complementarity determining regions (CDRs) (Figure 1B, arrows).

**Figure 1.**
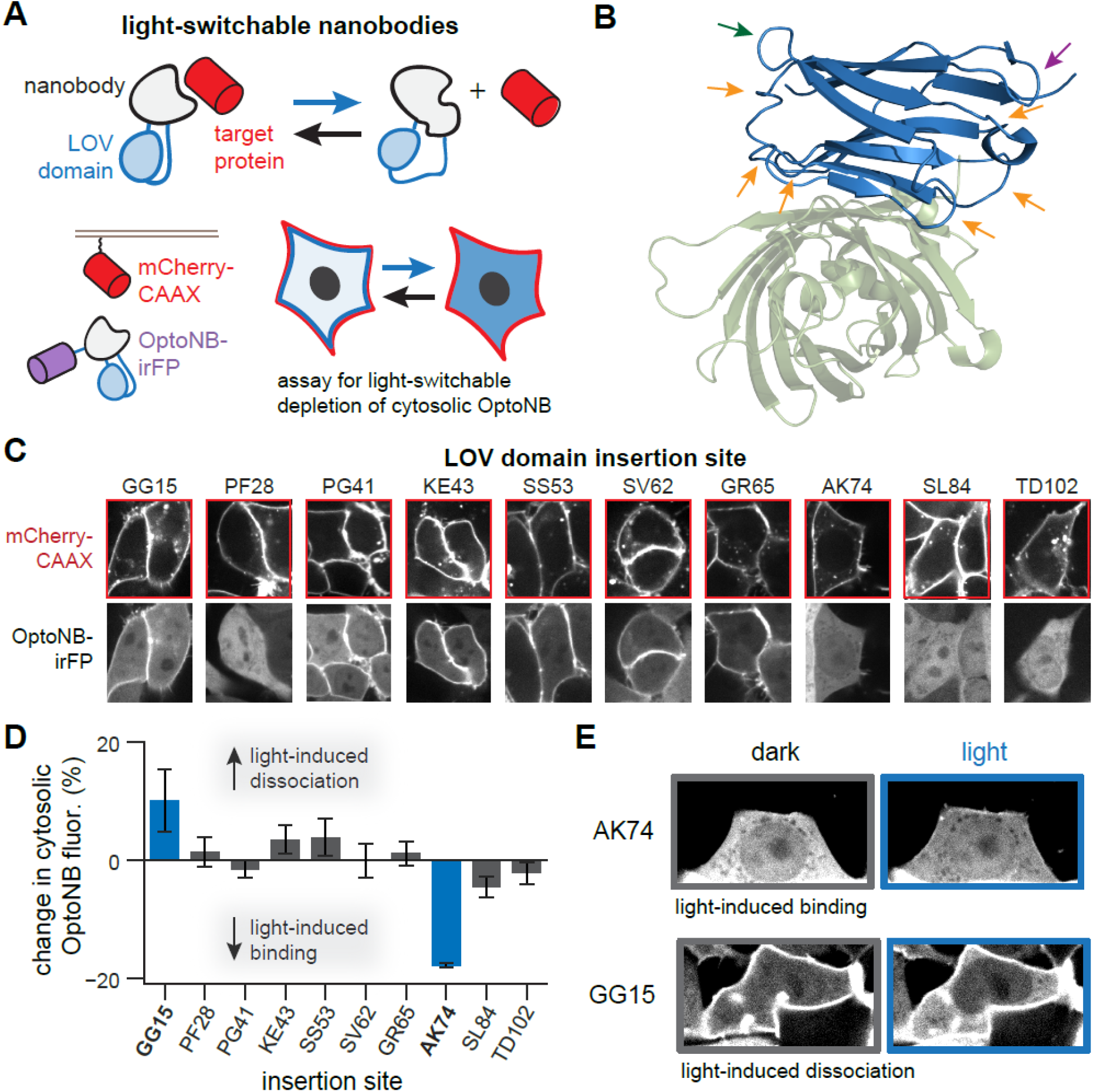
An initial screen for light-controllable nanobodies (OptoNBs). (**A**) Schematic of approach. By insertion into a solvent-exposed turn or loop, the light-switchable AsLOV2 domain (blue) could modulate the conformation of a nanobody (gray), thus allosterically altering its ability to bind to a target protein (red). Cytosolic irFP-fused OptoNBs were assayed for translocation to membrane-bound mCherry in the presence or absence of blue light. (**B**) Crystal structure of the LaG4 nanobody (blue) bound to EGFP (green), (PDB:3OGO), indicating loops targeted for LOV domain insertion (arrows). Two loops are highlighted containing Ala74 (green arrow) and Gly15 (purple arrow). (**C**) Results for insertion at 1-2 sites in all 8 loops. HEK293 cells expressing membrane-tethered mCherry (mCherry-CAAX) and cytosolic OptoNB-irFP (OptoNB) are shown, prior to blue light illumination. (**D**) Quantification of change in cytosolic intensity for at least 10 cells per OptoNB variant in **C**. An increase in cytosolic OptoNB fluorescence corresponds to light-induced dissociation from membrane-bound mCherry, and *vice versa* for light-induced decrease in cytosolic irFP. (**E**) Images before (gray box) and after (blue box) light stimulation for two OptoNB variants, LaM8-AK74 and LaM8-GG15, in HEK293T cells also expressing mCherry-CAAX, showing light-dependent changes in OptoNB localization in both cases.

We first set out to establish a cell-based assay for evaluating whether candidate opto-nanobodies exhibited light-switchable binding. Prior work has shown that monitoring cytosol-to-membrane translocation is a fast and sensitive method to characterize light-switchable binding (Figure 1A), revealing changes in protein localization for a diverse set of heterodimerization pairs and affinities^3,5,17^. We thus generated a set of HEK293 cell lines expressing a membrane-tagged mCherry target protein (mCherry-CAAX) and one of 10 candidate OptoNBs that were each fused on their C-terminus to an infrared fluorescent protein (OptoNB-irFP) (Figure 1A, **lower panel**, and 1C). We then imaged each cell line to determine the OptoNB’s subcellular localization in the presence or absence of 450 nm blue light.

The initial screen yielded diverse results for different nanobody-LOV fusions. Crucially, we observed redistribution between cytosol and membrane for two OptoNB fusions: those targeting the “GG15” and “AK74” insertion sites on Loops 1 and 6 of the nanobody (Figure 1B, green and purple arrows). (We name insertion sites based on the nanobody’s flanking amino acids and the position number of insertion, so AK74 corresponds to the sequence …-Ala-AsLOV2-Lys-… with AsLOV2 inserted after Ala74.) We found that light stimulation of these two sites triggered opposite effects on nanobody-mCherry binding: light-induced dissociation in the case of GG15 and binding in the case of AK74 (Figure 1D-E). This set of chimeras with opposite response to light was surprising, as prior studies that took advantage of LOV insertion reported only light-induced disruption of protein function, which was explained by the increased flexibility of the light-stimulated state disrupting the fusion protein’s active state^19,22^. In contrast, our data on the AK74 chimera suggests that the AsLOV2 dark state can also disrupt function, which is restored upon blue light illumination. We constructed a second round of cell lines to test additional insertion sites in Loops 1 and 6 where we obtained hits (positions 15-17 and 72-77) (**Figures S1A**). We observed similar light-induced changes at one additional site within each loop (GS16 and DN72), demonstrating that multiple sites within a single loop can be used to achieve photoswitchable binding control (**Figure S1B**).

Our initial results demonstrate that LOV-nanobody chimeric proteins are indeed capable of photoswitchable binding. They also reveal two distinct classes of optoNBs, those for which binding occurs in the light and others for which it occurs in the dark. The ability to induce binding by switching to dark conditions is rare, as most previously-developed optogenetic tools exhibit light-induced binding; yet the few existing light-suppressible optogenetic tools available^23,24^ have already proven useful for probing T cell signaling^25^, controlling metabolic flux^26,27^, and studying the consequences of protein phase separation^15^. Photoswitchable domain insertion thus holds promise for engineering light-based control of protein-protein interactions.

### An optimized LOV domain improves OptoNB function

Our initial screen also revealed that, for some optoNBs (GG15, NA73, and MG77), light unexpectedly triggered nuclear export (**Figure S1C-D**). Nuclear export observed even in cells that did not express membrane mCherry; furthermore, it was quickly reversed in the dark for the GG15 and MG77 variants but was irreversible for the NA73 variant. We thus next set out to improve the performance of our initial OptoNBs in two ways: eliminating undesired nuclear-cytosolic translocation of the nanobody and achieving a larger change in binding between dark and illuminated conditions.

We hypothesized that the light-induced nuclear/cytosolic translocation might arise due to light-triggered exposure of a nuclear export sequence (NES), as has been engineered in prior AsLOV2-based optogenetic tools^12,14^. Indeed, amino acid sequence analysis revealed a canonical NES (LxxxLxxLxL, where x is any amino acid and L is a hydrophobic amino acid that is often leucine) spanning the junction between the C-terminal Jα helix and nanobody for the GG15 and MG77 insertion sites (**Figure S1E**). (We did not observe a canonical NES for the NA73 variant, suggesting a different mechanism underlies its irreversible nuclear export.) We thus sought to truncate residues from the nanobody’s C-terminus to eliminate undesired NES activity. We also reasoned that truncating amino acids from the nanobody-AsLOV2 junction may have an additional benefit, enabling tighter conformational coupling between the LOV domain and nanobody. A close examination of the crystal structure of AsLOV2 (PDB: 2V0U) suggested that removing linker residues at both the N and C termini of the AsLOV2 domain could more tightly couple it to the nanobody (Figure 2A).

**Figure 2.**
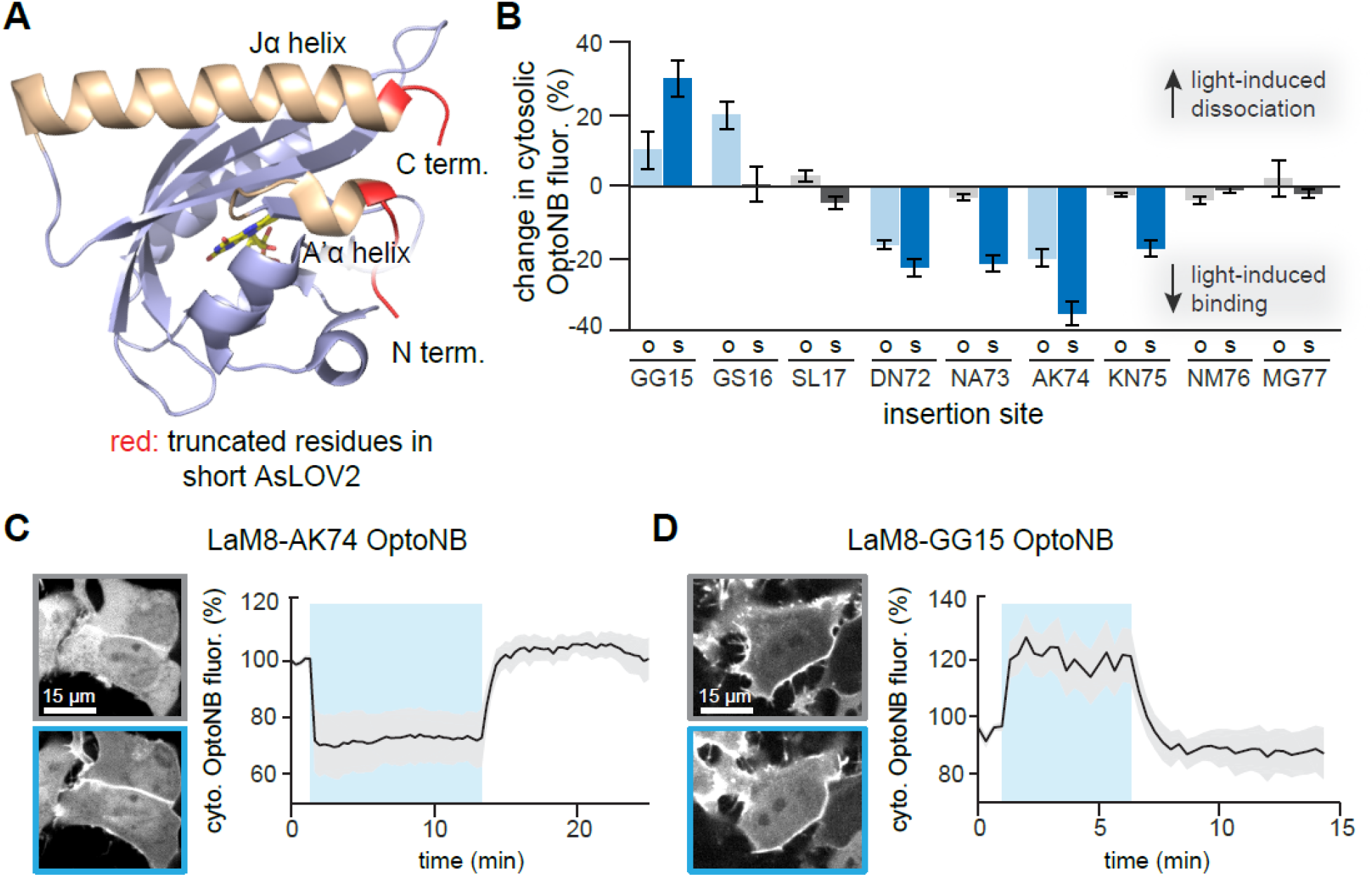
An optimized short LOV domain for OptoNB engineering. (**A**) AsLOV2 crystal structure (PDB:2V0U) indicating amino acids removed (red) to generate the optimized short AsLOV2 (408-543) for nanobody insertions. (**B**) Comparison of photoswitchable OptoNB binding in original AsLOV2 (‘o’) and short AsLOV2 (‘s’) for 9 insertion sites near the original two hits, GG15 and AK74. Data analyzed is steady-state change in cytosolic intensity for at least 5 cells per construct. (**C-D**) Light-induced membrane/cytosol translocation in HEK293T cells for the LaM8-AK74 (in **C**) and LaM8-GG15 (in **D**) OptoNBs. The percent change in cytosolic intensity from the original, dark-equilibrated value is shown. Curves and shaded regions indicate mean ± SD for at least 10 cells.

Based on this rationale, we constructed a ‘short LOV’ (sLOV) domain, comprising residues 408-543 of *Avena sativa* Phototropin 1 (versus residues 404-546 in Figure 1 and 404-547 in Ref 19), and re-screened insertion sites near our two initial hits (Figure 2B). We no longer observed light-dependent nuclear export in any sLOV insertions, consistent with the role of the C-terminal NES in this phenomenon. Additionally, light-induced binding changes were enhanced in 5 of 6 cases (GG15, DN72, NA73, AK74 and KN75) compared to the original AsLOV2 constructs (Figure 2B). We confirmed that light-switchable target binding could be reversibly toggled on and off for both light- and dark-inducible OptoNB variants by measuring localization in cycles of darkness and blue light illumination (Figure 2C-D; **Movies S1-2**). These results demonstrate that insertion of an optimized LOV domain can eliminate undesired nuclear/cytosolic translocation and generate opto-nanobodies with enhanced photoswitchable binding.

### OptoNBs can be constructed for multiple nanobodies and target proteins

Our initial OptoNB designs were in the context of a single binding pair: the LaM8 nanobody and its mCherry binding partner. We next sought to test whether light-induced binding or dissociation generalizes to other nanobodies or targets. Alignment of multiple nanobody sequences (our original LaM8 nanobody, the higher-affinity mCherry nanobody LaM4, and an anti-EGFP nanobody LaG9) indicates that both the AK74 and GG15 insertion sites are located in conserved regions that are distinct from the hypervariable complementarity-determining regions (CDRs) (Figure 3A)^21^. We thus hypothesized that similar effects would be elicited upon LOV domain insertion in the same sites of each nanobody.

**Figure 3.**
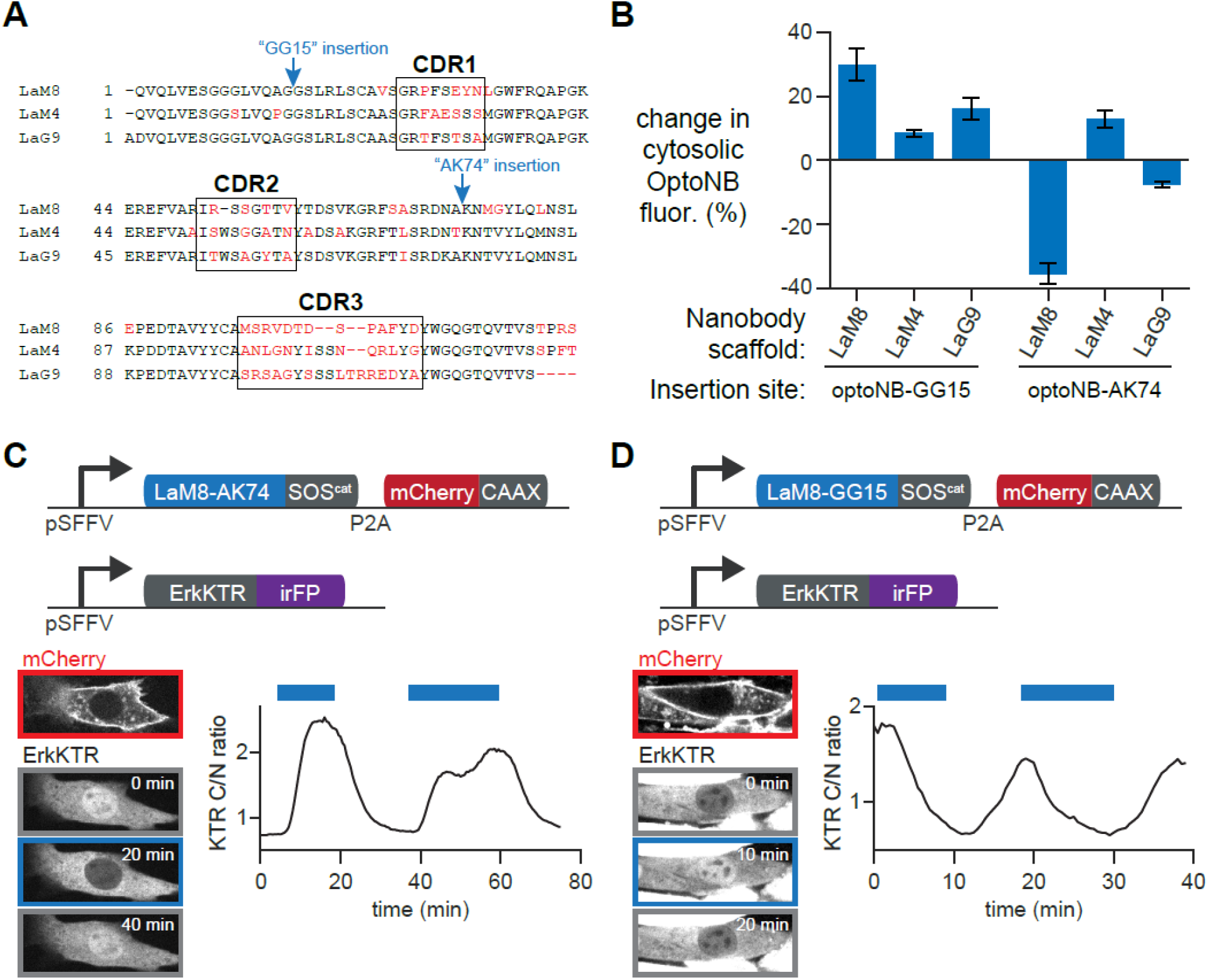
OptoNB design is generalizable and can be used to control intracellular signaling. (**A**) Sequence alignment for LaM8 (anti-mCherry), LaM4 (anti-mCherry), and LaG9 (anti-EGFP) nanobodies. Residues that are not conserved are shown in red and the three complementarity-determining regions (CDRs) are boxed. Blue arrows indicate AsLOV2 insertion sites. (**B**) Light-induced translocation of LaM8, LaM4 and LaG9 OptoNBs with LOV insertion at AK74 or GG15. The change in cytosolic intensity from the original, dark-equilibrated value is shown. Curves and shaded regions indicate mean ± SD for at least 5 cells. (**C-D**) NIH3T3 cell lines harboring OpoNB-controlled Ras/Erk pathway activity using LaM8-AK74 (in **C**) and LaM8-GG15 (in **D**). Upper diagrams show lentiviral constructs expressed in each cell: membrane-localized mCherry-CAAX, OptoNB-SOScat, and a live-cell biosensor of Erk activity (ErkKTR-irFP). Lower diagrams indicate mCherry and ErkKTR expression and localization for representative cells. Curves show the cytosolic to nuclear ratio of ErkKTR intensity for a representative cell during cycles of darkness and illumination (blue bars).

We generated sLOV insertions at the GG15 and AK74 positions in LaM4-irFP and LaG9-irFP fusion proteins. We then co-transduced cells with membrane mCherry or EGFP and monitored cytosolic OptoNB levels during cycles of blue light illumination or darkness (Figure 3B). We found that insertion at the GG15 position led to consistent light-inducible dissociation in all three nanobodies, although both LaM4 and LaG9 exhibited weaker dissociation than LaM8, possibly due to their higher-affinity target binding (0.18 and 3.5 nM, respectively, vs. 63 nM for LaM8)^21^. In contrast, AK74 insertion led to highly variable results across all three nanobodies: we observed light-induced binding in the cases of LaM8 and LaG9, but light-triggered dissociation for LaM4. Our results thus indicate that sLOV insertion at the same sites (GG15 and AK74) confers light-regulated target binding across multiple nanobodies (LaM8, LaM4, LaG9) and/or target proteins (GFP, mCherry). Insertion at the GG15 position appears to generally elicit light-triggered dissociation, whereas AK74 insertion appears to be more variable, with changes in both the sign and magnitude of photoswitchable binding observed between distinct nanobodies. This result is consistent with a model where light-induced destabilization of LOV domain contacts are more likely to disrupt protein function upon light stimulation than to restore it^19^, suggesting that the light-induced binding we observed in LaM8-AK74 may be a relatively unusual occurrence.

### Coupling OptoNBs to Ras/Erk signaling *in vivo*

Because our initial screen demonstrated light-directed control over protein binding in live mammalian cells, we reasoned that it should thus be possible to apply OptoNBs for light-based control over cellular functions without any further modification or optimization. To demonstrate this capability, we set out to construct an OptoNB-based variant of the OptoSOS optogenetic tool^28^. In this system, membrane localization of the catalytic domain of SOS (SOS^cat^) is used to trigger Ras activity^29^, activation of the Erk mitogen activated protein kinase cascade, and eventual responses including cell proliferation and differentiation. Light-induced signaling can also be easily visualized within minutes using the fluorescent Erk kinase translocation reporter (ErkKTR), a synthetic substrate that is exported from the nucleus to the cytosol within minutes of phosphorylation by active Erk^30^.

To reversibly trigger OptoSOS activation using nanobody-target binding, we generated NIH3T3 cell lines expressing an OptoNB-SOS^cat^ fusion protein (LaM8-AK74-SOS^cat^ or LaM8-GG15-SOS^cat^), membrane-localized mCherry (mCherry-CAAX) and an infrared fluorescent ErkKTR (ErkKTR-irFP) (Figure 3C-D; **Movies S3-4**). We found that Erk activity could be rapidly toggled on and off with each OptoNB variant (Figure 3C-D). As expected from our initial protein-binding results, Erk signaling has opposite responses depending on which OptoNB is used to recruit SOScat to the membrane, with light-induced activation in LaM8-AK74 OptoSOS cells (Figure 3C) and light-induced inactivation in LaM8-GG15 OptoSOS cells (Figure 3D). These results demonstrate that OptoNBs can indeed be deployed in cells to manipulate downstream cellular functions (e.g., Ras/Erk signaling) without further optimization.

### OptoNBs regulate protein binding *in vitro*

In addition to controlling intracellular protein-protein interactions, light-controlled nanobodies could be useful *in vitro* for a variety of applications, including extracellular reagents to modulate receptor-level responses^31^, and the ability to decorate light-switchable binders in biochemical purification columns to separate unmodified target proteins based on light stimuli ^32^. We thus set out to characterize the light-dependent performance of OptoNBs *in vitro* in a variety of assays: column-based separation, protein binding to OptoNB-coated agarose beads, and bio-layer interferometry-based measurement of OptoNB-protein binding kinetics.

As a first test of their function *in vitro*, we sought to test whether purified OptoNBs and their binding partners could be differentially separated in light and darkness using size exclusion chromatography. We expressed and purified the dark-inducible binder LaM4-TK74 and the light-inducible binder LaM8-AK74 from *E. coli* (see Supplementary Methods). We wrapped a Superdex 200 10/300 GE column with blue light-emitting diodes (LEDs; Figure 4A) or wrapped it in aluminum foil (to keep dark conditions), and flowed solutions containing OptoNB, mCherry, or both in a 1:1.2 molar ratio of NB to mCherry through the column (Figure 4B-C). We observed a strong light-dependent shift in retention time for both OptoNBs. In the case of LaM8-AK74, light-induced binding leads to a shorter retention time under illumination (Figure 4B, blue curve), and a longer complex retention time as well as a peak of free mCherry in the dark (Figure 4B, black curve; compare to red curve for free mCherry). LaM4-TK74 exhibits the converse response, with shorter retention in the dark and longer in the light, indicating light-induced dissociation as previously observed *in vivo* (Figure 4C).

**Figure 4.**
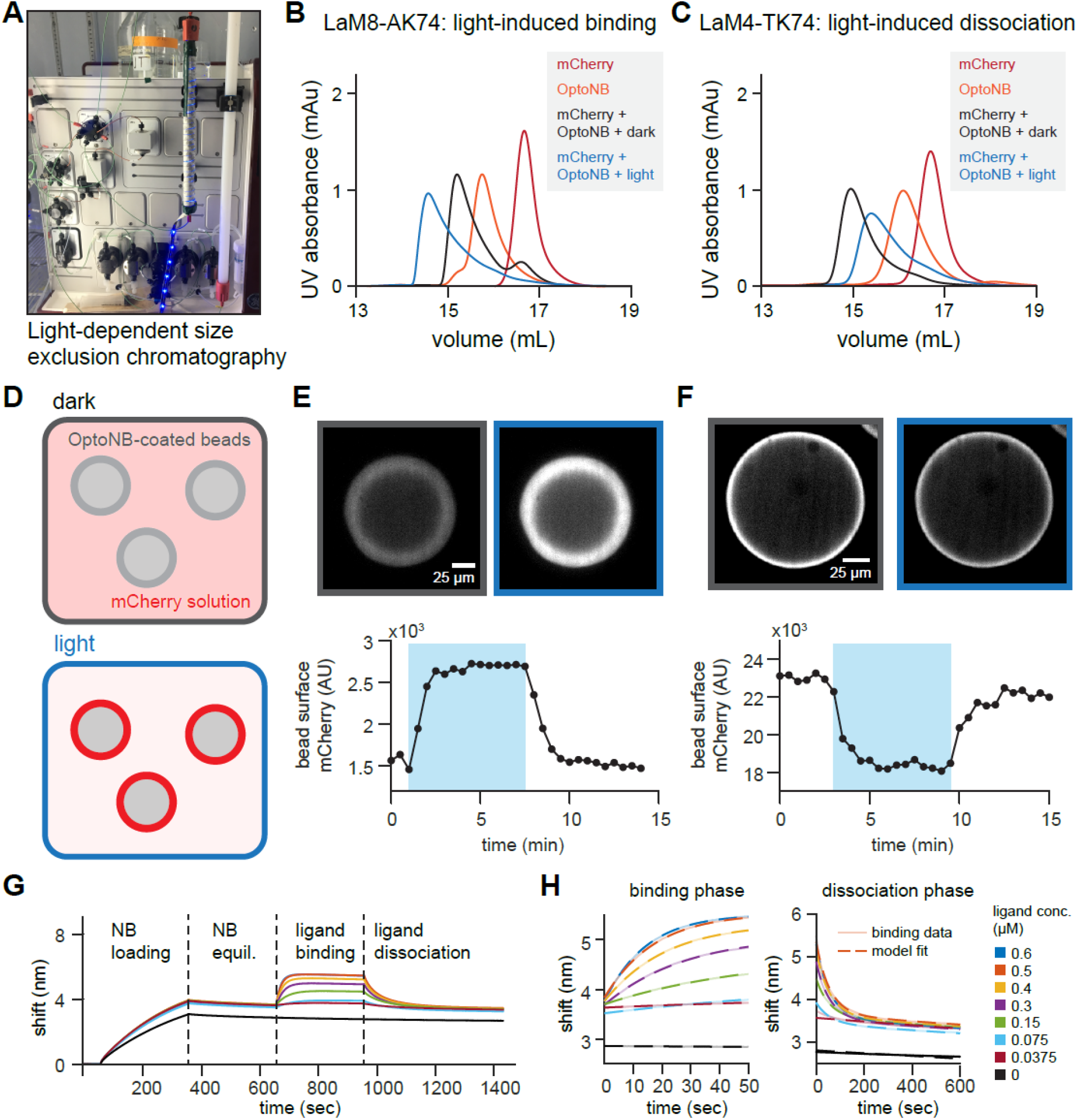
*In vitro* characterization of OptoNB binding. (**A**) Size exclusion chromatography (SEC) for light-dependent protein separation. The column was wrapped with 450 nm blue LEDs to allow for direct illumination of the protein during the SEC run, or in aluminum foil to keep it in darkness. (**B-C**) SEC elution profile for LaM8-AK74 (in **B**) and LaM4 TK74 (in **C**). Free mCherry, free OptoNB, and dark- and light-incubated OptoNB/mCherry mixtures are shown in the indicated curves. Shorter retention times indicate larger size and increased complex formation. (**D**) Schematic representing light-induced binding of mCherry to OptoNB-coated beads. His-tagged OptoNBs are immobilized on Ni-NTA agarose beads, while untagged mCherry is in the buffer solution surrounding the beads. A change in illumination conditions results in mCherry-OptoNB binding and a brighter bead surface in the mCherry channel. (**E-F**) Top panels: confocal mCherry images of beads coated with a mixture of His-tagged eGFP with His-tagged LaM8-AK74 in a 200:1 ratio (in **E**) or LaM8-GG15 in a 1000:1 ratio (in **F**). Beads were placed in 1 μM mCherry solution (in **E**) or 2 μM mCherry solution (in **F**). A 450 nm LED was toggled on and off (blue shading indicates LED illumination). Bottom panels: quantification of bead surface intensity during cycles of darkness and blue light illumination. (**G**) Representative bio-layer interferometry (BLI) traces for quantifying nanobody-protein binding and dissociation kinetics. Four phases indicate His-tagged OptoNB loading onto the Ni-NTA coated tips, equilibration in buffer, binding to different concentrations of soluble mCherry, and mCherry dissociation into buffer. (**H**) Raw data (solid lines) and best-fit traces simultaneously fit to simple mass-action kinetic binding model (dashed lines) for the Ni-NTA sensor – immobilized LaM8 nanobody binding to soluble mCherry at eight concentrations.

Many *in vitro* and extracellular applications of protein binders are based on interactions on surfaces (e.g., cell surface receptor binding; affinity-based purification using bead-tethered antibodies). We thus sought to test whether OptoNB-target interactions could be controlled on the surface of agarose beads (Figure 4D). We purified His-tagged OptoNBs (His_6_-LaM8-AK74 and His_6_-LaM8-GG15), as well as His_6_-GFP and His_6_-mCherry from *E. coli* (see Supplementary Methods). To obtain beads with different surface densities of immobilized OptoNBs, we incubated nickel-NTA-coated agarose beads with solutions containing different ratios of anti-mCherry His_6_-OptoNB and His_6_-GFP (where His_6_-GFP was used as a bead surface blocking agent) to achieve different surface densities of OptoNBs.

To test for light-dependent mCherry binding, we incubated the OptoNB-labeled beads with soluble mCherry (His_6_-cleaved) and imaged mCherry fluorescence during cycles of blue light illumination (Figure 4E-F; **Movies S5-6**). We observed a light-dependent shift in surface mCherry fluorescence as expected for AK74- and GG15-based OptoNBs. However, the time required to saturate the beads with bound nanobody upon light activation depended strongly on the density of target protein on the beads (**Figure S2**). Complete mCherry binding was achieved within 10 sec when beads were labeled with a 0.5%:99.5% LaM8-AK74:GFP protein ratio. In contrast, beads labeled with 100% LaM8-AK74 were only gradually saturated with mCherry over ~1 h. This phenomenon is likely due to local depletion of mCherry near the illuminated bead’s surface at high labeling densities, a picture that is consistent with the approximately linear increase in surface mCherry observed at high OptoNB concentrations^33^. In sum, we find that OptoNBs exhibit light-switchable binding to their targets across a broad range of contexts, from the mammalian intracellular environment to surface- and solution-based binding *in vitro*.

### Measurement of lit- and dark-state OptoNB binding kinetics

As a final characterization of OptoNB activity, we set out to perform quantitative measurements of their binding affinity and kinetics in the lit and dark states using bio-layer interferometry (BLI). We reasoned that BLI would be ideal for quantifying light-triggered protein-protein interactions due to its compatibility with sample illumination and accurate quantification of both binding kinetics and affinity. Binding measurements in the dark state can be made by taking advantage of the C450V point mutation in AsLOV2 which prevents photoadduct formation, rendering AsLOV2-based optogenetic tools light-insensitive^34,35^. Although a variety of lit state mutants have been characterized that destabilize docking of the C-terminal Jα and N-terminal A’α helices^36,37^, it is unclear whether these mutations fully mimic the lit state when these helices are constrained by being inserted into loops of a chimeric fusion, as in our OptoNBs.

We first expressed and purified His-tagged variants of the parental LaM8 nanobody, the LaM8-AK74 and LaM8-GG15 OptoNBs, and OptoNB mutants that are expected to be light-insensitive and locked in either the dark state (C450V equivalent) or lit state (I532E A536E equivalents)^38^. For each BLI run, one of these nanobody variants was loaded onto Ni-NTA-coated sensors, equilibrated in buffer, exposed to varying concentrations of mCherry to measure the association phase, and finally washed to measure the dissociation phase (Figure 4G). The binding curves at all mCherry concentrations were then globally fit to a simple mass-action chemical kinetic model of binding and dissociation, from which estimates of k_on_, k_off_, and K_D_ were obtained (Figure 4H, **Supplementary Code**). The global fitting procedure was able to fit the data well in each case (**Figure S3**).

The resulting kinetics and affinities are presented in Table 1. We measured an affinity of 230 nM for wild-type LaM8, which differed somewhat from the 63 nM affinity reported previously^21^, possibly due to differences in assay design and procedures used to fit binding curves. We found that the LaM8-GG15 OptoNB also exhibited sub-micromolar affinity for mCherry in its dark state (610 nM) that was weakened to 1.7-2.4 μM in the lit state. In contrast, the LaM8-AK74 variant exhibited weaker affinity for mCherry in its dark state (19.9 μM) than its lit state (3.65-3.79 μM), just as we had observed *in vivo* and *in vitro*. All lit state measurements agreed closely between illuminated, photosensitive OptoNBs and lit-state mutants, suggesting that these mutants accurately reflect the nanobody’s lit state. Finally, we note that in each case, the affinity change upon illumination was explained primarily by changes in the dissociation rate k_off_, with little change in the association rate k_on_. This observation is consistent with a light-dependent change in the complementarity of the nanobody’s binding site for its target protein, decreasing overall affinity by shortening the residence time of the bound complex. In sum, we demonstrate that bio-layer interferometry can be used to obtain binding kinetics and affinities for native lit-state optogenetic tools, without relying on mutants that may not perfectly approximate this state for a particular application. Applied to our LaM8-GG15 and LaM8-AK74 OptoNBs, BLI reveals a 3- to 5.5-fold change in binding affinity between lit and dark states *in vitro*.

**Table 1.** Binding kinetic measurements for lit- and dark-state OptoNBs using bio-layer interferometry (BLI). Columns list the nanobody variant used, illumination conditions, and best-fit kinetic rate constants to the raw BLI traces.

| Variant | Illumination | $k_{on}$ ($\mu\text{M}^{-1} \text{s}^{-1}$) | $k_{off}$ ($\text{s}^{-1}$) | $K_D$ ($\mu\text{M}$) |
| --- | --- | --- | --- | --- |
| LaM8 | None | 0.076 | 0.018 | 0.23 |
| LaM8-GG15 C450V (pseudo-dark) | None | 0.037 | 0.023 | 0.61 |
| LaM8-GG15 I532E A536E (pseudo-lit) | None | 0.042 | 0.102 | 2.41 |
| LaM8-GG15 | 450 nm light | 0.044 | 0.076 | 1.73 |
| LaM8-AK74 C450V (pseudo-dark) | None | 0.024 | 0.471 | 19.9 |
| LaM8-AK74 I532E A536E (pseudo-lit) | None | 0.036 | 0.13 | 3.65 |
| LaM8-AK74 | 450 nm light | 0.04 | 0.153 | 3.79 |

## Discussion

Here we have shown that a simple strategy for constructing light-sensitive proteins – the insertion of a photoswitchable domain into a target protein – can be used to produce light-sensitive nanobodies (OptoNBs). Nanobodies are a class of proteins with high potential as biological reagents for cell and developmental biology^39,40^, biotechnology^32^, and therapeutic applications^31,41^ because of their small size, ease of expression and purification in both bacterial and eukaryotic cells, and high-affinity binding to a growing list of target proteins. We show that OptoNBs can be produced from multiple nanobody scaffolds, show activity against multiple target proteins, can exhibit either light-inducible or light-dissociable responses, and can be functionally coupled to cell signaling *in vivo* and column-based assays *in vitro*. We further describe design variants to optimize OptoNB function, including an optimized, shortened AsLOV2 scaffold that improves photoswitchable binding in 5 of the 6 cases tested and eliminates a side pathway – light-switchable nuclear export – that could be relevant for intracellular applications.

The ability to control nanobody binding with light offers considerable promise for a variety of applications. Nanobodies can be raised against a broad range of target peptides through affinity maturation in immunized camelids, making it easy to envision proteins whose binding can be toggled on and off from a broad range of endogenous protein targets. In many cases, nanobody binding to an endogenous target protein could competitively inhibit the target protein’s normal function, opening the door to reversible, specific loss-of-function control. This is particularly intriguing, as most existing optogenetic tools regulate binding between two engineered domains, leaving endogenous processes unaffected. OptoNBs also have high potential for use as light-sensitive reagents outside of cells. Light-switchable binders could in principle be used to reversibly neutralize ligands or cell surface receptors on cells, tissues, or organisms that have not been genetically modified^31^; they could also enable separation of target proteins from complex mixtures without a need for affinity tags that may interfere with the target protein’s function^42^.

Importantly, we find that the existing mCherry-targeting LaM8 OptoNBs are already viable optogenetic tools for intracellular applications. LaM8-AK74 exhibits photoswitchable mCherry binding that is comparable to other high-quality optogenetic tools, with low basal binding in the dark and large-scale protein translocation to the membrane upon illumination. We also demonstrate that this localization change can be harnessed for regulating intracellular processes by constructing OptoNB-based variants of the OptoSOS system for Ras/Erk pathway control. It should be noted that mCherry is frequently used in intracellular labeling experiments, so LaM8-based OptoNBs could be immediately deployed as a ‘backwards-compatible’ strategy for altering the intracellular localization of these mCherry-tagged proteins in the cell lines or organisms where they have already been developed.

Nevertheless, further improvements are likely to be needed before OptoNBs reach their full potential, especially in the case of *in vitro* applications. Our analysis of binding affinities in the lit and dark states indicates that our LaM8-based OptoNBs exhibit up to a 5.5-fold change between lit and dark states, spanning an overall 30-fold range of affinities between the tightest (610 nM for LaM8-GG15’s dark state) and weakest binders (19.9 μM for LaM8-AK74’s lit state). These results indicate that there is still substantial room for improvement of photoswitchable OptoNB function. Prior studies have designed engineered LOV domains and cognate peptides with up to 50-fold changes in affinity between lit and dark states^4^. Moreover, in an accompanying study targeting a monobody against the Abl SH2 domain, we report that sLOV insertion chimeras can elicit a ~100-fold change in affinity (see accompanying manuscript: Carrasco-Lopez et al).

How might OptoNB function be further improved? One possible route to improvement could be to improve the allosteric coupling between the LOV domain’s insertion site and nanobody’s target-binding surface. A larger change in binding affinity may thus be achieved by testing additional nanobody insertion sites using the shortened LOV domain sequence or screening additional LOV domain variants with tighter dark-state binding^43^. We also note the apparent discrepancy between the large localization change observed in cells (Figure 2C) and the *in vitro* binding assay (Table 1). It is thus possible that OptoNBs respond differently in these different biochemical environments. For example, nanobodies contain a disulfide bond that may be differentially formed depending on the reducing environment and might alter the nanobody’s sensitivity to allosteric control. OptoNBs thus hold considerable promise for delivering programmable, light-controlled binding for broad range of applications inside and outside cells.

## Materials and Methods

### Plasmid Construction

DNA containing LaM8, LaM4, and LaG9 was kindly gifted by Professor Kole Roybal (UCSF). All DNA was cloned using backbone PCR and inFusion (Clontech). pHR vectors were used for mammalian cell experiments and pBAD vectors were used for bacterial protein overexpression. AsLOV2 404-546 and AsLOV2 408-543 were ordered as gene blocks from IDT and used to insert into the nanobodies using inFusion (Clontech). Any additional plasmids were constructed using backbone or insert PCR and inFusion (Clontech). Stellar competent cells were transformed by all the plasmids for amplification and DNA storage.

### Lentivirus Production and Transduction

HEK 293T cells were plated on a 6 or 12 well plate and grown up to 40% confluency. At that point they were co-transfected with desired pHR plasmid and lentiviral packaging plasmids (pMD and CMV) using Fugene HD (Promega). Virus was collected after approximately 48 hours, filtered using 0.45 mm filter and 2 μL of polybrene and 40 μL of HEPES were added to the viral particles. Either HEK 293T or NIH 3T3 cells were plated on a 6-well plate and infected with 200-500 μL of the virus at 40% confluency. Viral media was replaced by growth media 24 hours post infection and imaging was done at least 48 hours post the infection time. Media used for all cell culture maintenance contained DMEM, 10% FBS, penicillin, and streptomycin.

### Cell imaging

For imaging, 0.17 mm glass-bottomed, black-walled 96-well plates (In Vitro Scientific) were used. Glass was first treated with 10 μg/mL of fibronectin in PBS for 20 min. Cells were then plated and allowed to adhere onto the plate. 50 μL of mineral oil was added on top of the media prior to imaging to limit media evaporation. For RAS/Erk signaling experiments, cells were switched to starvation media prior to imaging (plain DMEM + 20 mM HEPES buffer, with no added serum). Cells were washed 3 times with starvation media and then equilibrated in starvation media for at least 3 hours prior to imaging.

The mammalian cells were kept at 37°C with 5% CO_2_ for the duration of all imaging experiments. Imaging was done using Nikon Eclipse Ti microscope with a Prior linear motorized stage, a Yokogawa CSU-X1 spinning disk, an Agilent laser line module containing 405, 488, 561 and 650 nm lasers, an iXon DU897 EMCCD camera, and a 40X oil immersion objective lens. A 450nm LED light source was used for photoexcitation with blue light, which was delivered through a Polygon400 digital micro-mirror device (DMD; Mightex Systems).

### Protein Expression

All proteins were expressed using pBAD N-His vector. Nanobody and opto-nanobody plasmids were transformed into Shuffle T7 Express *E.coli* cells (NEB) and eGFP and mCherry plasmids were transformed into One Shot Top 10 cells (Invitrogen). A single colony was used to inoculate a 10 mL 2x YT overnight culture supplemented with 200 μg/mL of Carbenicillin (Carb). The following day the culture was used to inoculate 0.5 L of 2x YT/Carb media that was shaken at 37 °C and 250 rpm until it reached an OD600 of approximately 1.0. Subsequently, the temperature was decreased to 20 °C and protein expression was induced by adding 0.2% Arabinose. The culture was shaken in the dark for approximately 18 hours followed by harvesting the cells by centrifugation at 4 °C and 12000g. If the protein was not purified right away, the pellets were stored at −80 °C.

For the purification, the 0.5 L cell pellet was thawed and resuspended in 25 mL of resuspension buffer (50 mM Tris pH 8.0, 150 mM NaCl) with 0.4 mM phenylmethanesulfphonylfluoride (PMSF) as well as a tablet of cOmplete Mini (Roche) and 14 μL of β-mercapthoethanol. Cells were lysed using a sonicator and the supernatant clarified by centrifugation at 250,000 x g for 1 hour. Subsequently, FMN (0.25 mg/mL) was added to the supernatant with ~ 30 min incubation to ensure a homogenous distribution of the chromophore. 3-4 mL of Ni-NTA superflow resin (Qiagen) were loaded onto a column and equilibrated with the resuspension buffer. The supernatant was loaded onto the column followed by 100 mL washes with resuspension buffer containing increasing concentrations of imidazole of 10, 20, and 30 mM. The protein was eluted at 250 mM imidazole and dialyzed overnight against resuspension buffer with the protein purity determined by SDS-PAGE. Protein concentrations were determined by recording the Abs_280_ and the following extinction coefficients for eGFP, mCherry, LaM8, and OptoLaM8NBs, 24,995 M^−1^ cm^−1^, 34,380 M^−1^ cm^−1^, 24,535 M^−1^ cm^−1^, and 47,905 M^−1^ cm^−1^, respectively.

### Size exclusion chromatography

The size exclusion chromatography was performed on an AKTA Pure system (GE Healthcare) at 4°C. The Superdex 200 Increase 16/300 GL column (GE Healthcare) was equilibrated with 50 mM Tris pH 8.0, 150 mM NaCl and this buffer was used for all subsequent SEC experiments. The purified proteins were assembled in 1:1.2 molar ratio of mCherry:nanobody or mCherry:OptoNB. The final volume of the proteins loaded onto the column was 50-200 μL, depending on the protein concentration. For experiments run in the dark, the lights were turned off, the chromatography refrigerator was covered in a black blanket, and the column was wrapped in aluminum foil. For experiments run in the light the column was wrapped with blue LED string (Grainger). Before loading the proteins onto the column, the mixed samples were incubated for 20 min at room temperature, either in blue light or dark, according to the experiment that was being performed.

### Agarose Bead Imaging

Ni-NTA agarose resin (Qiagen) was first equilibrated with 50 mM Tris pH 8.0, 150 mM NaCl buffer. To competitively label the resin beads, solutions of 500 μL 1:10, 1:100, and 1:1000 nanobody:eGFP solution was loaded onto 200 μL of resin slurry with the excess protein washed away with the same buffer. 50 μL of 1 μM mCherry (with its His-tag cleaved off using TEV protease) was added onto 0.17 mm glass-bottomed black walled 96 well plate (In Vitro Scientific). 2 μL of the nanobody bead slurry was added to the well with mCherry solution and incubated for at least an hour and up to overnight. The same microscope setup (imaging and blue light excitation) was used to image the beads as previously described for the cell imaging, except for the use of a 20X air objective lens for the beads.

### Bio-layer interferometry measurements of binding kinetics

Measurements for the on rates (k_on_), off rates (k_off_), and affinity constants (K_D_) for LaM8, LaM8 AK74, and LaM8 GG15 nanobodies were performed on Octet RED96e instruments (ForteBio). Ni-NTA sensors (ForteBio) were first equilibrated in 50 mM Tris pH 8.0, 150 mM NaCl buffer for 10 min prior the measurement. Clear 96-well plates were used for the measurements and wells were filled with 200 μL of buffer or sample. During the experimental run the sensors were first immersed in a buffer to record the baseline, then switched to load the His-tagged nanobody onto the sensor and back into the buffer to remove unbound nanobody. To measure the association rate the sensors were subsequently moved into a well with 8 different concentrations of tagless mCherry including a control with 0 mM mCherry. To measure the k_off_ the sensors were then moved into wells with a buffer and the dissociation rate was recorded. In order to measure binding kinetics of the light state, the lid to the Octet remained open during the measurement and a blue LED panel was held above the 96-well plate keeping the protein in the light state for the duration of the experiment. The raw binding and unbinding data were simultaneously fit to models of the binding and unbinding reactions:

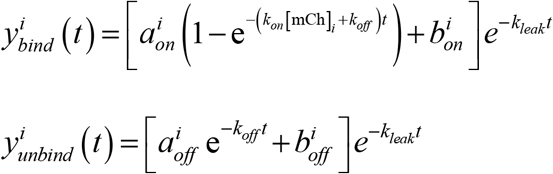

This model incorporates the following dependent and independent variables:
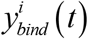 refers to the i^th^ binding curve
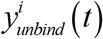 the i^th^ unbinding curve
[mCh]_*i*_ refers to the concentration of mCherry used for the i^th^ binding curve
*t* is the time elapsed since the start of the binding/unbinding phase.

It also includes the following parameters

*k*_*on*_ is the on-rate (same across all binding and unbinding curves)

*k*_*off*_ is the off-rate (same across all binding and unbinding curves)

*k*_*leak*_ represents the slow unbinding of His-tagged OptoNB from the probe, leading to a gradual decay of signal.

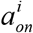 is the total change in signal due to mCherry binding for the i^th^ curve

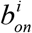 is the signal baseline during the binding phase

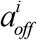 is the total change in signal due to mCherry unbinding for the i^th^ curve

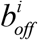 is the signal baseline during the unbinding phase

The model thus contains 4\**n* + 3 parameters, where *n* is the number of distinct mCherry concentrations tested. Fits were performed using nonlinear gradient descent using the MATLAB `fmincon` function. The MATLAB code used to perform the fits is included as Supplementary Code.

## Author contributions

Conceptualization, A.A.G., J.L.A., and J.E.T.; Methodology, A.A.G., E.M.Z., M.Z.W., and J.E.T.; Investigation, A.A.G., E.M.Z., M.Z.W., A.G.G., and C.C.L.; Writing – Original Draft, A.A.G. and J.E.T.; Writing – Review & Editing, all authors; Funding Acquisition, A.A.G., J.L.A. and J.E.T.; Supervision, J.L.A and J.E.T.

## Acknowledgements

We thank all members of the Toettcher and Avalos labs for helpful comments, and Nicole Neville for her initial opto-nanobody studies. We also thank the Biophysics Core Facility and Venu Vandavasi for help with the bio-layer interferometry measurements. This work was supported by NIH grant DP2EB024247 (to J.E.T.), the Pew Charitable Trusts, the U.S. DOE Office of Biological and Environmental Research, Genomic Science Program Award DESC0019363, NSF CAREER Award CBET-1751840, and a Camille Dreyfus Teacher-Scholar Award (to J.L.A.), NIH Fellowship F32GM128304 (to A.A.G.) and NIH Fellowship F30CA206408 (to A.G.G.).

