## Supplementary Text for "Optogenetic control of protein binding using light-switchable nanobodies"

**This PDF file includes:**

Supplementary Figures 1-3
Supplementary Video 1-4 Legends

### Supplementary Figures and Legends

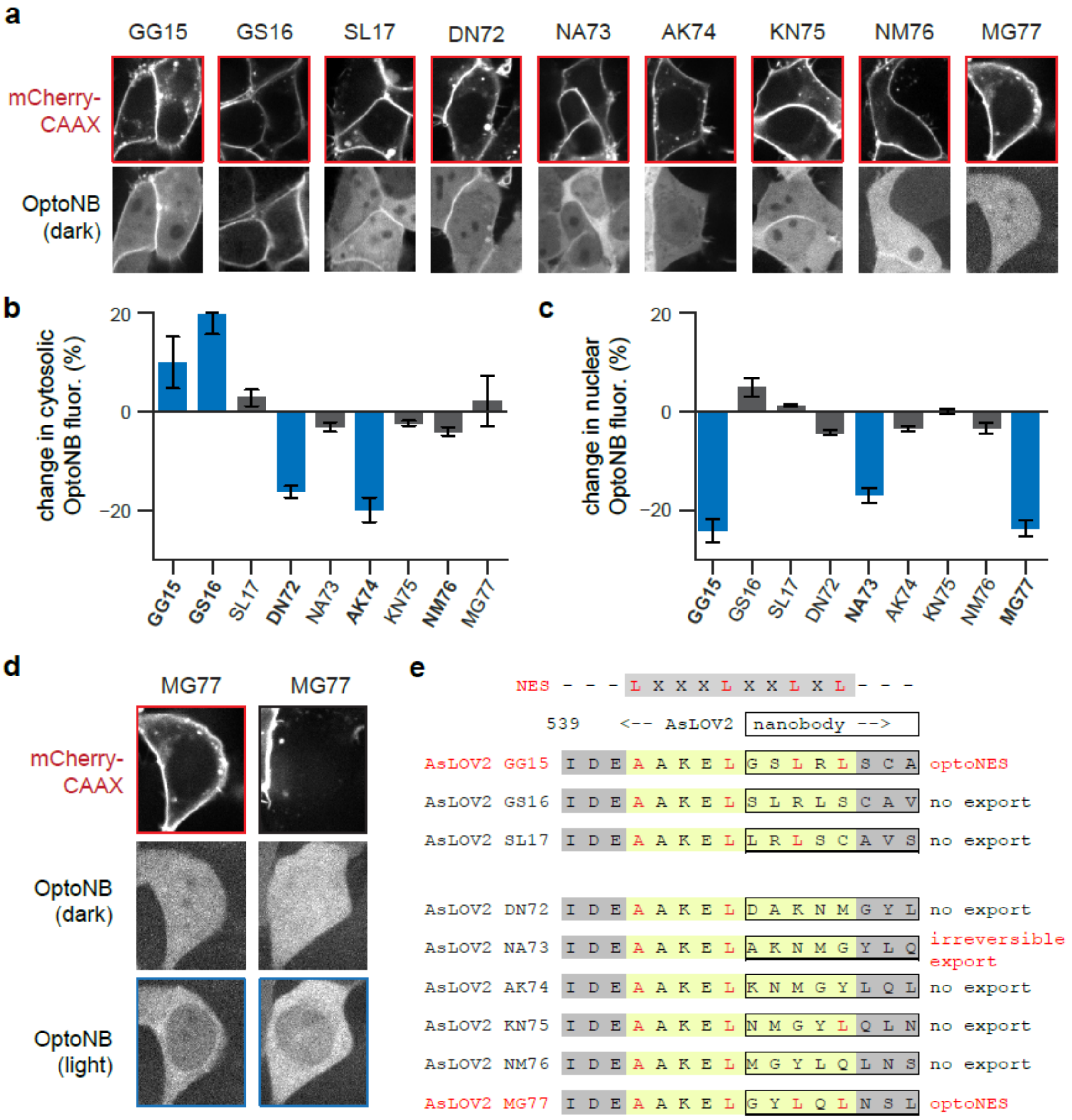


**Figure S1. Membrane-cytosolic and nuclear-cytosolic translocation of initial OptoNBs.** (**a**) OptoNB variants around GG15 and AK74 insertions. Upper: mCherry CAAX expression on the membrane. Lower: Initial OptoNB-iRFP localization in the dark. (**b**) Change in cytosolic fluorescence of OptoNB variants around GG15 and AK74 insertions indicating light induced dissociation for GG15 and GS16, and light induced binding for DN72 and AK74. (**c**) Change in nuclear fluorescence of OptoNB in cells without mCherry CAXX membrane component. Reversible light induced nuclear export is observed in GG15, and MG77, along with light induced irreversible nuclear export in NA73. (**d**) Light-induced nuclear export in MG77 variant. Left panels: a cell expressing LaM8-MG77 and mCherry-CAAX; right panels show a cell expressing only the LaM8-MG77. Nuclear translocation is observed in both cases. (**e**) A canonical nuclear export sequence (LxxxLxxLxL) is mapped onto the sequence at the C-terminal junction between AsLOV2 and the nanobody fusion for various OptoNB insertion sites. Light-induced nuclear translocation is marked in red text.

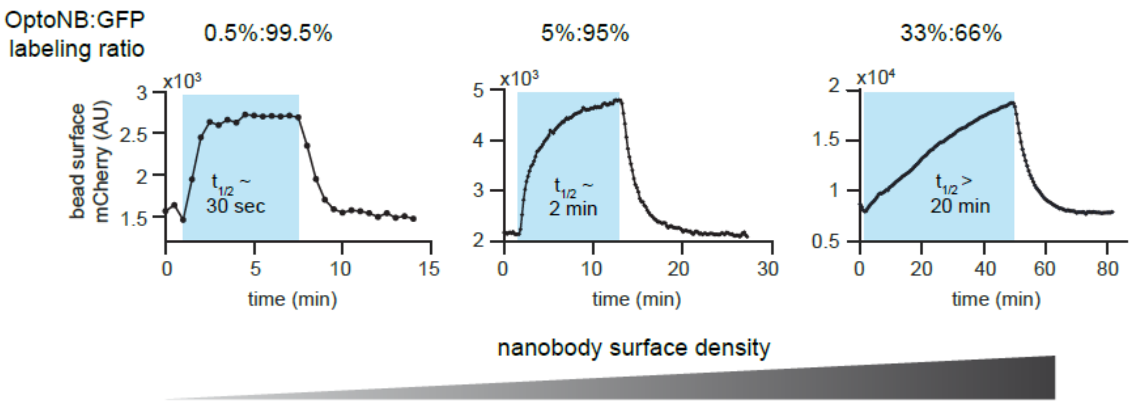


**Figure S2. Quantification of mCherry binding to LaM8-AK74 coated beads.** His-tagged LaM8-AK74 OptoNBs and His-tagged eGFP were immobilized on the surface of NiNTA agarose beads in various ratios (0.5%:99.5%; 5%:95%; 33%:66%). Each was incubated in a solution of 1 μM mCherry, and mCherry intensity on the bead surface was quantified over time. Increasing the nanobody surface density resulted in slower mCherry binding/unbinding kinetics as well as a shift from exponential to linear kinetics, as predicted from the formation of a diffusion-limited mCherry depletion layer in the vicinity of the bead.

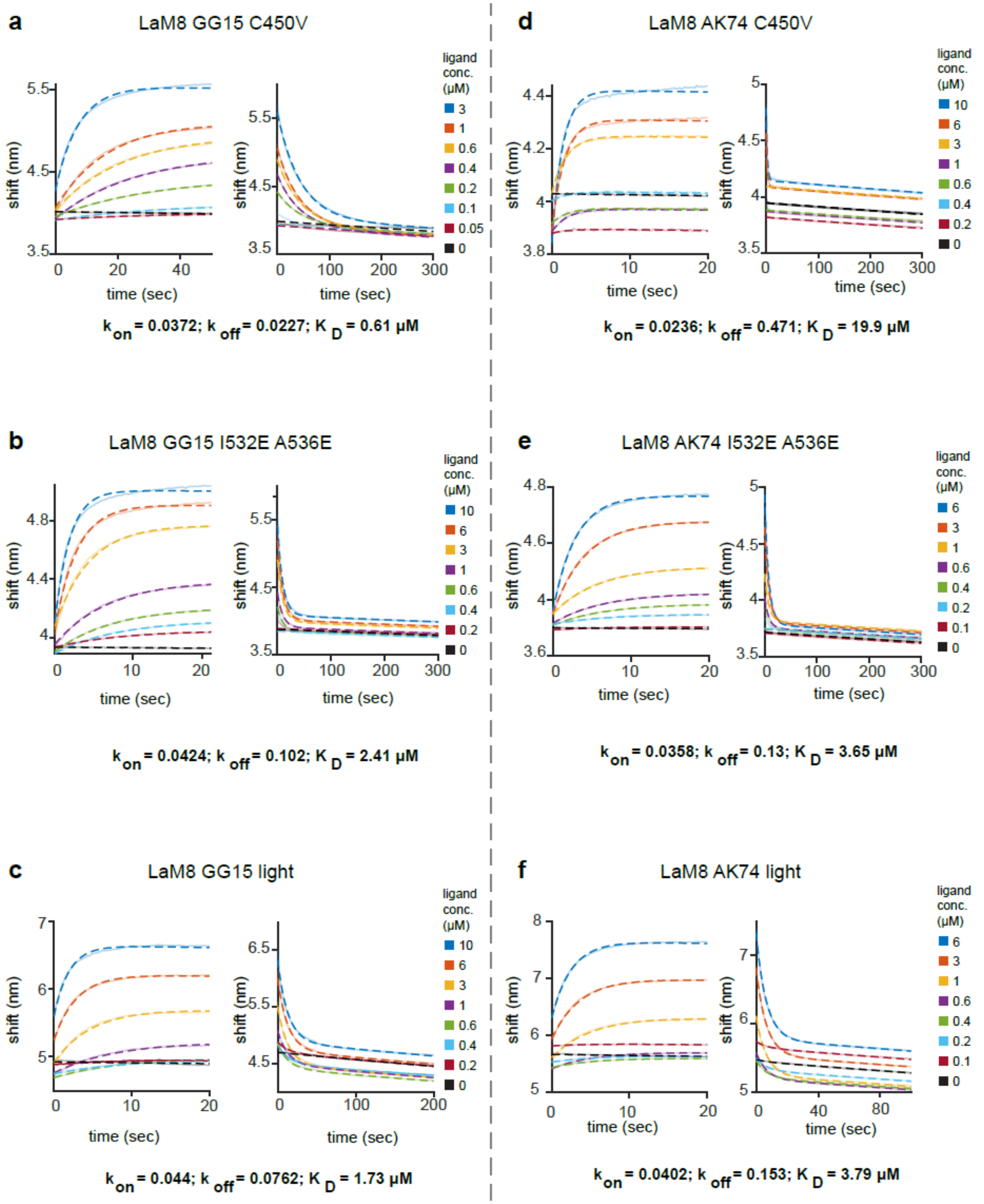


**Figure S3. Bio-layer interferometry (BLI) traces for LaM8 GG15 (a-c) and LaM8 AK74 (d-f).** All plots show the raw BLI shifts for tip-immobilized OptoNB variants over time in various concentrations of mCherry. Raw data are shown as solid curves; model fits are shown in dashed curves. In order to measure the binding constants from the dark/preilluminated state, we used a dark-like C450V mutant that is unable to transduce light absorption into a conformational change (in **a** and **d**). For lit-state measurements we compared lit-like I532E/A536E double mutant (in **b** and **e**) to a true lit-state produced by illuminating the BLI sensors with a constant 450 nm LED light input (in **c** and **f**). The reported $k_{on}$ values are all in $\mu M^{-1} s^{-1}$, $k_{off}$ values are in $s^{-1}$, and all $K_D$ values are in μM.

### Supplementary Code

One MATLAB script (`scriptRunOctetFit.m`), two MATLAB functions (`construct_kinetics_fit.m` and `run_fits.m`), and one sample data file (`RawData0.log`) are supplied in accompanying files. To use, unzip all four files to a single directory and, in MATLAB, run `>>scriptRunOctetFit`. The code will automatically identify the binding and unbinding phases and fit a single set of global parameters to obtain estimates of $k_{on}$, $k_{off}$ and $K_d$. To run on additional user-defined data, the only required parameters are the location of a new raw dataset and the ligand concentrations that were used for each tip.

### Supplementary Video Legends

**Movie S1.** Time-lapse imaging of HEK293T cells expressing the LaM8-AK74_irFP OptoNB (shown) and membrane-localized mCherry-CAAX (not shown). Cells were imaged using a 60X oil objective every 20 sec for 30 min as 450 nm blue LED illumination was toggled on and off (indicated by the blue box in the lower-left corner). Timer indicates mm:ss; scale bar indicates 10 μm.

**Movie S2.** Time-lapse imaging of HEK293T cells expressing the LaM8-GG15_irFP OptoNB (shown) and membrane-localized mCherry-CAAX (not shown). Cells were imaged using a 60X oil objective every 20 sec for 30 min as 450 nm blue LED illumination was toggled on and off (indicated by the blue box in the lower-left corner). Timer indicates mm:ss; scale bar indicates 10 μm.

**Movie S3.** Time-lapse imaging of an NIH3T3 cell expressing LaM8-AK74_SOS$^{cat}$ OptoNB fusion protein (not shown), membrane-localized mCherry-CAAX (not shown), and ErkKTR-irFP (shown). Cells were imaged using a 60X oil objective every 30 sec for 80 min as 450 nm blue LED illumination was toggled on and off (indicated by the blue box in the lower-left corner). Timer indicates hh:mm; scale bar indicates 10 μm.

**Movie S4.** Time-lapse imaging of an NIH3T3 cell expressing LaM8-GG15_SOS$^{cat}$ OptoNB fusion protein (not shown), membrane-localized mCherry-CAAX (not shown), and ErkKTR-irFP (shown). Cells were imaged using a 60X oil objective every 30 sec for 80 min as 450 nm blue LED illumination was toggled on and off (indicated by the blue box in the lower-left corner). Timer indicates hh:mm; scale bar indicates 10 μm.

**Movie S5.** Time-lapse imaging of a NiNTA-coated agarose bead with His-tagged LaM8-AK74 and His-tagged GFP immobilized on its surface in a 5%:95% ratio, in a solution of 1 μM untagged, soluble mCherry. The bead was imaged using a 20X air objective every 30 sec for 80 min as 450 nm blue LED illumination was toggled on and off (indicated by the blue box in the lower-left corner). Timer indicates hh:mm; scale bar indicates 50 μm.

**Movie S6.** Time-lapse imaging of a NiNTA-coated agarose bead with His-tagged LaM8-GG15 and His-tagged GFP immobilized on its surface in a 0.1%:99.9% ratio, in a solution of 2 μM untagged, soluble mCherry. The bead was imaged using a 20X air objective every 30 sec for 30 min as 450 nm blue LED illumination was toggled on and off (indicated by the blue box in the lower-left corner). Timer indicates hh:mm; scale bar indicates 50 μm.
