## Supplementary Code for "Optogenetic control of protein binding using light-switchable nanobodies": construct_kinetics_fit.m

```
function fcn = construct_kinetics_fit(conc, t1, x1, t2, x2, toPlot)
if nargin < 6 || isempty(toPlot)
    toPlot = 0;
end

t1 = t1-t1(1);
t2 = t2-t2(1);

x1m = zeros(size(x1));
x2m = zeros(size(x2));

fcn = @fit_fcn;

    function SSE = fit_fcn(p)
        ka  = p(1);
        kd  = p(2);
        k2  = p(3);
        
        
        % Load blocks of 4 parameters for each set of data
        c = 0;
        while 1
            try
            c = c + 1;
            aon(c)  = p(4*(c-1)+4);
            bon(c)  = p(4*(c-1)+5);
            aoff(c) = p(4*(c-1)+6);
            boff(c) = p(4*(c-1)+7);
            catch, break, end
        end
        
        x1m = zeros(size(x1));
        x2m = zeros(size(x2));
        
        for i = 1:size(x1,2)
            % THIS LINE IS RIGHT!!!! ->>>
            x1m(:,i) = (aon(i)*(1-exp(-(ka*conc(i) + kd)*t1)) + bon(i)).*exp(-k2*t1);
            % THIS LINE IS WRONG!!! STUPID JARED.
%             x1m(:,i) = (aon(i)*(1-exp(-(ka*conc(i)*t1))) + bon(i)).*exp(-k2*t1);
        end
        
        for i = 1:size(x2,2)
            x2m(:,i) = (aoff(i)*exp(-kd*t2) + boff(i)).*exp(-k2*t2);
        end
        
        SSE = sum((x1m(:)-x1(:)).^2) + sum((x2m(:)-x2(:)).^2);
        
        if toPlot
            figure(2)
            clf
            subplot(2,1,1)
            plot(t1, x1)
            hold on
            ax = gca;
            ax.ColorOrderIndex = 1;
            plot(t1, x1m, '--')
            title(sprintf('k_{on} = %0.3g; k_{off} = %0.3g; K_D = %0.3g', p(1), p(2), p(2)/p(1)))
            subplot(2,1,2)
            plot(t2, x2)
            hold on
            ax = gca;
            ax.ColorOrderIndex = 1;
            plot(t2, x2m, '--')
            SSE = 0;
%             keyboard
            drawnow
%             pause
        end
    end
end
```
