## Supplementary Code for "Optogenetic control of protein binding using light-switchable nanobodies": run_fits.m

```
function pfit = run_fits(fpath, p)
% Assume that the file to load is always RawData0.log
fname = 'RawData0.log';

%% Load data from the RawData0.log file

% Import data
tmp = importdata([fpath '\' fname]);

%% Extract raw Octet data
% The first column corresponds to time points
t = tmp.data(:,1);
% All subsequent columns correspond to data traces
x = tmp.data(:,2:2:end);

% Truncate to remove NaN entries
x = x(:,any(x));

% Do some basic noise filtering to smoothen noisy trajectories using the
% h-minima transform
for i = 1:size(x,2)
    xh = imhmin(x, p.hmin);
end

%% Find switches between conditions using the derivatives of the traces

% Takes advantage of the fact that big jumps happen when solutions are
% changed!

xm = max(abs(diff(xh)),[],2);
[v I] = sort(xm, 'descend');

% Don't get fooled by big derivative times that are too close to
% even bigger ones nearby:
for k = 1:20 % remove nearby ones
    for m = 1:50
        ii = find(I(k+1:end) == I(k)+m);
        I(k+ii) = nan;
    end
end
I = I(~isnan(I));

%% Plot all the phases to verify that they were correctly identified

i_s = sort(I(1:4))+1;
ts = t(i_s);

figure(1),clf
set(gcf, 'position', [0 420 662 190])
plot(t, x)
for i = 1:length(ts)
    line([1 1]*ts(i), [0 10])
end
title(p.outname, 'interpreter', 'none')
pause

%% Plot the association and dissociation phases to verify that they were correctly identified

t1  = t(i_s(3):i_s(4)-1);
x1  = x(i_s(3):i_s(4)-1,:);
i1 = find(t1 <= t1(1) + p.maxt1);
t1 = t1(i1);
x1 = x1(i1,:);
t2 = t(i_s(4):end);
x2 = x(i_s(4):end,:);
t12 = [t1; t2];
x12 = [x1; x2];

figure(2),clf
set(gcf, 'position', [0 0 535 541])
subplot(3,1,1)
plot(t1, x1)
subplot(3,1,2)
plot(t2, x2)
subplot(3,1,3)
plot(t12, x12)
pause

%% Construct the functions to fit to these data traces!

F = construct_kinetics_fit(p.conc, t1, x1, t2, x2);
Fplot = construct_kinetics_fit(p.conc, t1, x1, t2, x2, 1);

%% Get the vector of a, b, kon, koff, k2 initial parameter guesses

ka0   = 1e-2;
kd0   = 1e-1; 
k20   = 3e-4;
p0 = [ka0 kd0 k20];
for i = 1:size(x1,2)
    bon0  = x1(1,i);
    aon0  = (x1(end,i)-bon0)*exp(k20*(t1(end)-t1(1)));
    boff0 = x2(end,i)*exp(k20*(t2(end)-t2(1)));
    aoff0 = x2(1,i)-boff0;
    p0 = [p0 aon0 bon0 aoff0 boff0];
end
p0(p0 < 0) = 0;

Fplot(p0)

%% Perform the fit!

options = optimoptions('fmincon');
options.Display = 'iter';
options.MaxFunctionEvaluations = 1e4;
options.OutputFcn = @(p,a,b) Fplot(p);

pfit = fmincon(F, p0, [], [], [], [], p0/1e2, p0*1e2, [], options);

%% Plot the fit results

figure(3),clf
set(gcf, 'position', [0 23 535 541])
Fplot(pfit)

%% Save file output
saveas(gcf, [p.outname], 'fig');
```
