## Supplementary Code for "Optogenetic control of protein binding using light-switchable nanobodies": scriptRunOctetFit.m

```
fpath = pwd; % point to the path where the "RawData0.xls" file is located

% Set the output filename
p.outname = 'TEST FIT';

% the mCherry concentrations used in each lane of the instrument (in micromolar)
p.conc = [0.6 0.5 0.4 0.3 0.150 0.075 0.0375 0];

% Maximum time period to use during the association phase.  It is important
% to truncate the association phase so that it is not dominated by
% the steady-state phase
p.maxt1 = 80; 

% This is a smoothing parameter that does not play a substantial role
% except in the case of VERY weak BLI signals (not for any of our published 
% data)
p.hmin = 0.1; 

% Run the fitting procedure!
run_fits(fpath, p)
```
